# The low-field effect in radical pairs: a zero-field singlet-triplet basis picture

**DOI:** 10.64898/2026.04.05.716627

**Authors:** J. R. Woodward

**Author notes:** Electronic mail.

## Abstract

We present a new formulation of the low-field effect (LFE) in spin-correlated radical pairs based on a zero-field singlet-triplet basis for the isotropic spin Hamiltonian. The aim is to provide a description that is both formally rigorous and mechanistically transparent, especially in the regime of weak magnetic fields such as the geomagnetic field. For the standard model radical pair containing a single spin 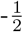 nucleus, we show that the conventional singlet-triplet basis obscures the distinct dynamical roles of the hyperfine and Zeeman interactions. In the zero-field S-T basis, by contrast, the mechanism separates cleanly: isotropic hyperfine coupling mixes singlet-doublet and triplet-doublet states, whereas the weak-field Zeeman interaction mixes triplet-quartet and triplet-doublet states without directly introducing an additional singlet-triplet coupling. The LFE is therefore revealed as a sequential process in which a weak field unlocks access from a triplet-only manifold to a singlet-accessible triplet manifold, from which hyperfine-driven singlet-triplet interconversion can occur. We then generalize this picture to radical pairs with arbitrary isotropic hyperfine structures by identifying maximal, interior, and, when present, minimal triplet-only manifolds in the zero-field spectrum. Finally, we introduce a practical blockwise dark-state recruitment measure for the triplet-only zero-field state space made singlet-accessible by a weak field, and show how this quantity depends on hyperfine symmetry, including the effects of equivalent nuclei. The resulting framework provides both a simple physical picture of the LFE and a general route to estimating its structural upper bound for arbitrary radical pairs.

## I. INTRODUCTION

The radical pair mechanism (RPM) is one of the leading proposals for how weak magnetic fields can influence chemistry and, potentially, biology. Since the 1970s it has been established that chemical reactions proceeding through radical-pair (RP) intermediates can show rates and yields that depend on applied magnetic fields^1–3^. More recently, the RPM has acquired wider significance through its proposed role in magnetoreception and cryptochrome photochemistry^4–8^. These developments have sharpened the need for descriptions of RP spin dynamics that are not only formally correct, but also mechanistically transparent across the very different field regimes relevant to spin chemistry and biology.

This challenge is especially acute for the low-field effect (LFE), namely the difference between RP behaviour in true zero magnetic field and in a very weak applied field such as the geomagnetic field. Early qualitative pictures were successful in explaining the contrast between zero-field and strong-field behaviour^2^. By contrast, the difference between true zero field and weak fields in the geomagnetic range is still much harder to summarize in comparably simple terms, even though it has been examined in detail in several important spin-dynamical studies^3,9,10^. This weak-field regime is also the one most directly implicated in biological and biomimetic discussions of the RPM, from chemical-compass models to cryptochrome photocycles and hypomagnetic effects^4–7,11^. At the same time, there is still no simple general method to estimate, from the hyperfine structure alone, the largest LFE that a given radical pair could in principle display.

The purpose of this work is therefore twofold. First, we seek a representation of RP singlet-triplet mixing in zero and weak magnetic fields that is both formally precise and conceptually accessible, especially to readers whose background is primarily biological rather than physical. Second, we use that representation to develop a structural measure of the maximum LFE available to an arbitrary RP on the basis of its hyperfine structure. The immediate motivation for pursuing such a reformulation came from our earlier analysis of triplet-born radical pairs, which highlighted both the importance of the LFE and the difficulty of describing it in a compact mechanistic language^12^. Both aims are achieved by changing the basis in which the RP spin states are described. As in many areas of quantum mechanics, the choice of basis is not merely a technical convenience: it can be the key step in revealing the mechanism of the underlying physics.

## II. BACKGROUND

### A. Core ingredients of the radical pair mechanism

The RPM explains how chemical reactions that proceed through the formation of short-lived RP intermediates can exhibit rates and yields that are altered by weak magnetic fields. A radical pair contains two radicals, each of which has one unpaired electron and therefore an electron spin quantum number 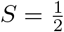 . The total spin state of the electron pair can therefore be either a singlet (*S* = 0) or a triplet (*S* = 1).

The magnetic-field sensitivity of RP reactions follows from two basic ingredients.

First, RP reactivity is spin selective. In most molecules the ground state is a spin singlet, because electrons pair in the lowest-energy orbitals in accordance with the Pauli principle. Consequently, when two radicals are close enough to react, they usually do so efficiently only when their total electronic spin state is singlet. In this common situation, singlet radical pairs are chemically reactive whereas triplet radical pairs are not. There are important exceptions (for example the ground state of molecular oxygen is triplet), but the singlet-reactive/triplet-unreactive picture captures the standard RPM setting.

Second, the mixing of RP spin states is magnetic-field sensitive. Once the radicals become sufficiently separated in space, their direct orbital overlap becomes negligible and the electron exchange interaction becomes small compared with the interactions of the individual electron spins with local spin-active nuclei. These hyperfine interactions drive coherent interconversion between singlet and triplet spin states. As a result, a triplet RP can become a singlet RP and vice versa. Since reaction is spin selective, the rate and efficiency of singlet-triplet mixing directly influence chemical yields and reaction rates. A magnetic field modifies this spin mixing and thereby modifies the chemistry.

This is the core of the RPM and can already be understood at an intuitive level without explicit resort to density matrices, reaction superoperators, or detailed time evolution under the spin Hamiltonian. In strong fields, relatively simple energy arguments are usually sufficient to explain which singlet and triplet states can and cannot mix^2^. In weak and zero magnetic fields, however, the standard state descriptions are much less effective. Even the clearest existing treatments require detailed quantum-mechanical state tracking and do not easily condense into a picture that a non-specialist can carry forward^3,9,10,13^. Our strategy will therefore be to begin with the familiar strong-field description, identify why it fails to give a transparent account of the LFE, and then replace it with a basis that does.

### B. The conventional singlet-triplet basis and the strong-field picture

The most direct description of the electron spins in a radical pair begins with the two individual electron spin states, which we denote by |*α*⟩ and |*β* ⟩. For two radicals, the product basis states are

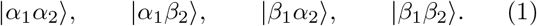

These basis states are convenient when the radicals are well separated and the two electron spins behave approximately independently.

For the RPM, however, the most important property of the RP is not the orientation of each individual electron but the total electronic spin character of the pair, because chemical reactivity distinguishes singlet and triplet states. The natural basis is therefore the set of eigenstates of the isotropic electron exchange operator (which are also the approximate eigenstates in a strong applied magnetic field), which are

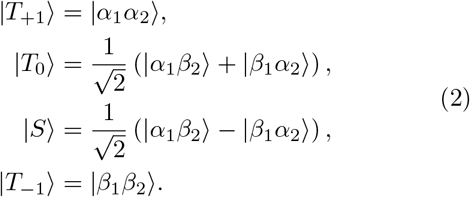

We refer to these as the conventional singlet-triplet or *S*–*T* basis states.

Consider first the simplest possible RP, containing just the two electrons and no nuclei. If both electrons experience the same Zeeman interaction, the spin Hamiltonian is

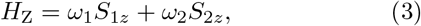

where *ω*_1_ = *ω*_2_ = *ω* = *gµ*_*B*_*B* and we set *ħ* = 1. In this case both the product basis states and the *S*–*T* basis states are eigenstates of the Hamiltonian. The magnetic field shifts the energies of |*T*_+1_ ⟩and |*T*_*−*1_⟩ relative to |*T*_0_⟩ and |*S* ⟩, but does not itself mix any of these states.

There are two ways in which singlet-triplet mixing can then arise. The first is the Δ*g* mechanism, in which the two radicals experience different local Zeeman interactions, so that *ω*_1_ ≠ *ω*_2_. In that case the Zeeman Hamiltonian develops the off-diagonal matrix element

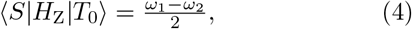

and coherent *S*–*T*_0_ mixing becomes possible. Because Δ*g* values are typically small, this mechanism is usually significant only in strong magnetic fields.

The second, and usually more important, source of singlet-triplet mixing is the hyperfine coupling of the electron spins to local spin-active nuclei. To see this in the simplest possible form, consider a single spin 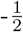 nucleus coupled to electron 1 with isotropic hyperfine coupling *a*. The hyperfine interaction is

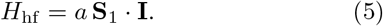

If the nuclear spin states are denoted by |*α*_*n*_⟩ and |*β*_*n*_⟩, the conventional *S*–*T* basis expands to the eight states

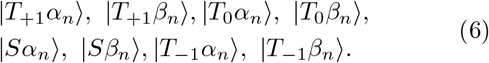

In this basis the Hamiltonian shows three kinds of mixing channel:

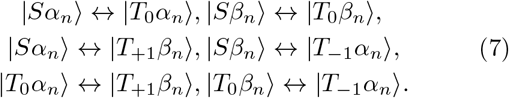

At high field, however, the *T*_+1_ and *T*_*−*1_ states move far away in energy from the *S* and *T*_0_ states. As a result, *S*– *T*_*±*1_ and *T*_0_–*T*_*±*1_ mixing become inefficient, leaving only the *S*–*T*_0_ channel active. This is the familiar strong-field picture: in zero field, all three triplet components are involved, whereas in a strong field the dynamics reduce effectively to *S*–*T*_0_ mixing.

This conventional picture is both useful and intuitive in the strong-field limit. The difficulty is that it becomes far less transparent when one attempts to compare true zero field with very weak field. In that regime the labels *S, T*_+1_, *T*_0_, and *T*_*−*1_ no longer cleanly isolate the physically relevant zero-field and weak-field dynamical processes.

## III. REVISITING THE LOW-FIELD EFFECT

### A. Why the conventional basis becomes opaque in weak field

The LFE is commonly analysed by working with the zero-field eigenstates of the same one-nucleus Hamiltonian discussed above. That procedure is entirely correct, but in the conventional *S*–*T* basis it remains conceptually awkward. In zero and weak magnetic fields the total electron spin *S* and its projection *m*_*S*_ are no longer good quantum numbers. As a result, singlet and triplet character are already mixed within the natural zero-field eigenstates, and the corresponding labels do not cleanly separate the hyperfine-driven and Zeeman-driven parts of the dynamics.

Previous discussions of the LFE have usually been framed either in the uncoupled product basis or in the conventional electron singlet-triplet basis, and have often described the phenomenon as a magnetic-field-induced modification of singlet-triplet interconversion^2,3,6,14^. These formulations have been indispensable for calculating reaction yields and for identifying the field range in which the effect occurs. In particular, the standard strong-field picture and the later weak-field treatments established the kinetic and spectroscopic language in which the problem is usually discussed^2,3,15^. A major step toward a sharper weak-field interpretation was then provided by Lewis *et al*.^9^. Working within the conventional singlet-triplet language, but analysing the exact zero-field eigenstates together with the projector matrix elements that govern the time-dependent singlet and triplet probabilities, they showed that lifting specific zero-field degeneracies introduces new time-dependent contributions in the *S* and *T*_0_ channels, rather than in *T*_*±*1_ as had previously been argued on angular-momentum grounds^16^. However, because both the hyperfine and Zeeman terms still appear as off-diagonal couplings in those representations, the distinct dynamical roles of the two interactions are not isolated as clearly as one might wish. The degeneracy structure responsible for the weak-field response is therefore less transparent than in a basis adapted to the zero-field Hamiltonian, and it becomes easy to speak loosely as if the applied field itself were directly generating the relevant singlet-triplet mixing.

What is needed, therefore, is a refactorisation of the state space based on the good quantum numbers in zero field, with the weak magnetic field introduced only afterwards as a perturbation. In such a basis the hyperfine and Zeeman interactions should, as far as possible, appear as distinct couplings between distinct classes of state. The purpose of the present reformulation is not to replace the earlier treatments, including the important analysis of Lewis *et al*., but to restate the mechanism in a basis adapted to the zero-field Hamiltonian. Their approach identifies which projection-operator channels acquire new time dependence when degeneracies are lifted; ours is designed to make the coupling topology itself explicit. In doing so it makes the angular-momentum structure transparent, shows which states are triplet-only and which carry singlet character, and separates hyperfine-driven mixing from Zeeman-driven mixing as cleanly as possible. This is the basis change that reveals the LFE most clearly and that also generalises naturally to arbitrary isotropic hyperfine networks.

### B. The zero-field S-T basis for one spin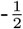 nucleus

We begin again with the single spin 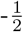 nucleus coupled isotropically to radical 1. In zero field, when only isotropic interactions are present, the good quantum numbers are the total spin on radical 1 and the total spin of the full spin system. We denote these by *F* and *J*, respectively:

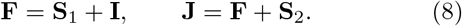

For this system we identify eight basis states. Four belong to the total-spin quartet manifold 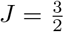 and are pure electron-triplet states. We denote this subset by *T*_*Q*_:

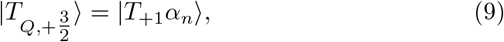

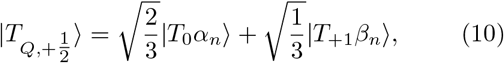

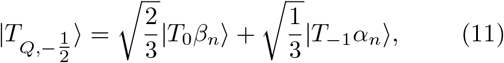

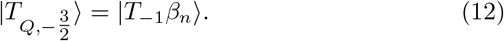

The remaining four states belong to the doublet manifold 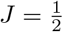. Two are pure electron singlets,

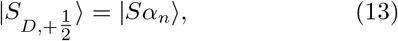

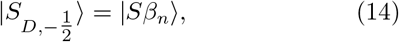

and two are pure electron triplets orthogonal to the *T*_*Q*_ states,

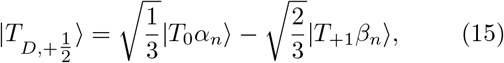

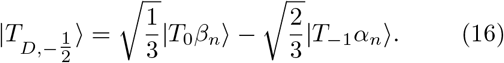

Thus, as in the conventional basis, there are six pure triplet states and two pure singlet states. The difference is that these states are now organised by the good zero-field quantum numbers and therefore reveal the weak-field dynamics much more cleanly. Fig 1. shows the 8 basis states for a one-proton RP in the three different basis state descriptions.

**FIG. 1.**
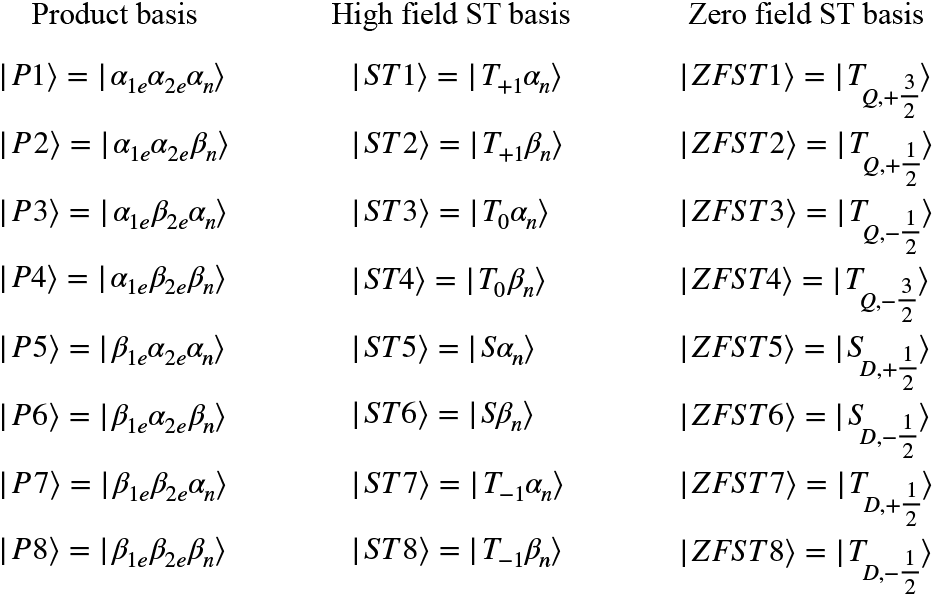
The eight basis states for a radical pair with s single spin 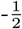 nucleus expressed in the product basis, the (conven-tional) high field singlet-triplet basis and the zero-field singlet-triplet basis introduced in this work.

### C. Zero-field mixing in the zero-field S-T basis

In zero field the Hamiltonian contains only the isotropic hyperfine interaction,

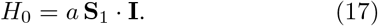

Its eigenstates are naturally labelled by |*F, J, M*⟩. The *F* = 1 manifold has energy 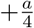 and is sixfold degenerate, while the *F* = 0 manifold has energy 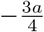 and is twofold degenerate. In terms of the zero-field S-T basis these states are

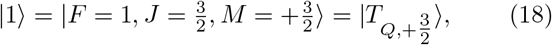

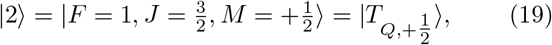

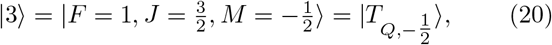

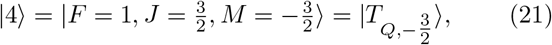

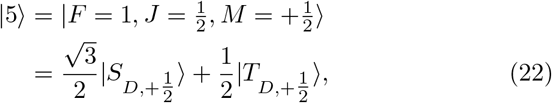

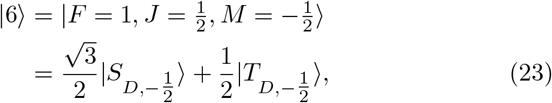

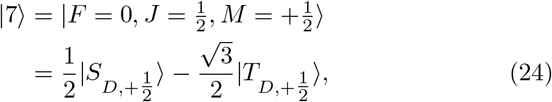

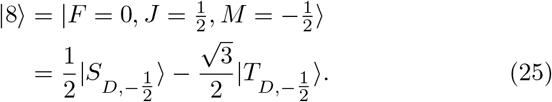

These states make the zero-field situation explicit. The *T*_*Q*_ states |1⟩ to |4⟩ are pure electron triplets. By contrast, the total-spin doublet states |5⟩ to |8⟩ are simple binary mixtures of pure electron singlet and triplet states. In other words, the zero-field singlet-triplet dynamics reside entirely in the doublet sector.

Ignoring the two stretched end states 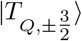, which cannot mix with anything else, the zero-field Hamiltonian in the basis

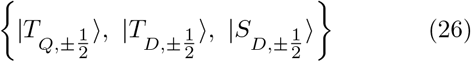

has the same form for 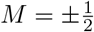 and is given by

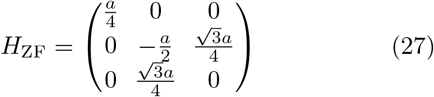

up to the obvious relabelling of the basis states. The key point is that the only nonzero off-diagonal coupling is the hyperfine-driven *T*_*D*_–*S*_*D*_ coupling, while the *T*_*Q*_ state remains uncoupled. Zero-field singlet-triplet mixing is therefore simply

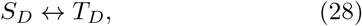

with no direct coupling to the *T*_*Q*_ sector. This already provides a much clearer picture than the conventional *S*–*T* basis.

### D. Weak-field mixing in the zero-field S-T basis

When a weak magnetic field is added, the electron Zeeman term must be expressed in the same basis. In the same 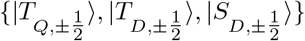 basis, the Zeeman Hamiltonian becomes

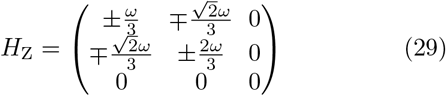

in the basis 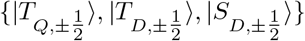. The crucial point is immediate: the Zeeman term mixes *T*_*Q*_ with *T*_*D*_, but does not mix *S*_*D*_ directly with either one.

The enhanced mixing in weak field is therefore not caused by the magnetic field directly turning on a new singlet-triplet matrix element. Instead, the magnetic field mixes the *T*_*Q*_ triplet states with the *T*_*D*_ triplet states, and the latter are already hyperfine-coupled to the singlet states. In schematic form,

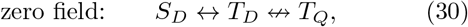

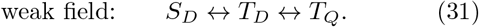

This is the central mechanistic result of the zero-field S-T basis. The LFE is not caused by the weak field directly inducing additional *S*–*T* mixing. Rather, it arises because the field unlocks additional triplet-triplet mixing, and this in turn enlarges the part of spin space from which hyperfine-driven singlet-triplet interconversion can proceed.

This resolves a common ambiguity in earlier qualitative descriptions. For equal electron *g* values, the field does not by itself generate a direct singlet-triplet matrix element; instead, it unlocks additional triplet-derived states that hyperfine coupling can then connect to the singlet-containing sector.

The zero-field S-T basis also gives a simple interpretation of the conclusion of Lewis *et al*. that the weak-field effect appears in the *S* and *T*_0_ channels. In the one-proton problem, the *T*_*D*_ states that are directly coupled to *S*_*D*_ contain one-third *T*_0_ character and two-thirds character in the corresponding *T*_*±*1_ component, whereas the recruited *T*_*Q*_ states have the opposite composition.

Weak-field mixing of *T*_*Q*_ into the singlet-accessible triplet sector therefore increases the *T*_0_ content of the triplet-derived states connected to *S*_*D*_. Thus the present mechanism is consistent with an apparent enhancement of *S*-*T*_0_ mixing in the conventional basis, while showing that the field acts indirectly through triplet-triplet recruitment rather than through a direct singlet-triplet coupling.

### E. A first abstraction of the mechanism

The zero-field S-T basis immediately allows a simple abstraction. There are three important subsets of states: quartet triplet states (*T*_*Q*_), doublet triplet states (*T*_*D*_), and doublet singlet states (*S*_*D*_). In zero field, only the *S*_*D*_ ↔*T*_*D*_ channel is active. In weak field, an additional *T*_*Q*_ ↔*T*_*D*_ channel becomes active. Thus the difference between zero field and weak field is not that singlet-triplet mixing suddenly appears, but that the weak field opens additional access to the triplet subspace that already communicates with the singlet sector.

This picture generalises naturally to more realistic radical pairs containing many nuclei. The one-nucleus *T*_*Q*_ manifold is replaced by a maximal triplet-only manifold *T*_max_, the triplet-doublet manifold *T*_*D*_ by an interior triplet sector *T*_int_, and the same sequential mechanism persists in the form

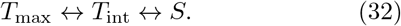

## IV. GENERALISATION TO ARBITRARY ISOTROPIC HYPERFINE COUPLINGS

We now generalise the preceding discussion to a radical pair containing arbitrary sets of isotropically coupled nuclei on the two radicals. The spin Hamiltonian is

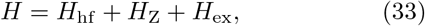

with

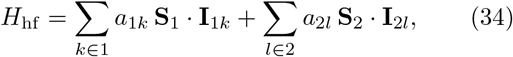

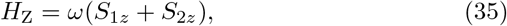

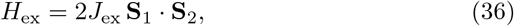

where the exchange term may be omitted when it is not required. Nuclear Zeeman, anisotropic hyperfine, dipolar, and Δ*g* terms are neglected.

For isotropic interactions the zero-field Hamiltonian remains rotationally invariant, so the exact zero-field eigenstates may still be classified by the total spin *J* of the full spin system and its projection *M* . To expose the angular-momentum structure, we define the total electron spin

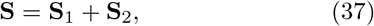

with *S* = 0 or 1, and the total nuclear spin

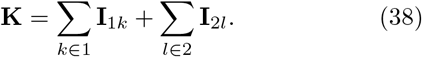

Coupling **S** and **K** then gives the total spin

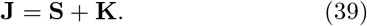

For a given total nuclear spin *K*, singlet-derived states (*S* = 0) occur only at

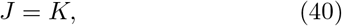

whereas triplet-derived states (*S* = 1) occur at

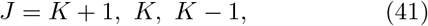

subject to the usual angular-momentum constraints. It follows that the largest possible total spin,

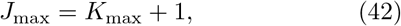

can only arise from electron-triplet states. Here

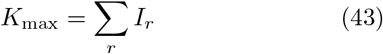

is the maximum allowed total nuclear spin, where the sum runs over all nuclei in the radical pair. We therefore define the manifold with *J* = *J*_max_ as the maximal triplet manifold, denoted *T*_max_. This is the direct many-spin generalisation of the one-nucleus *T*_*Q*_ manifold: it is always triplet-only and contains no singlet character.

The minimum total nuclear spin determines whether there is also a lower extremal triplet-only manifold. Let

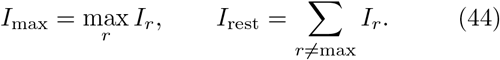

Then the minimum allowed total nuclear spin is

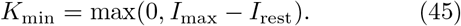

A minimal triplet-only manifold exists if and only if

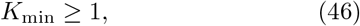

in which case its total spin is

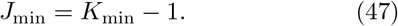

We denote this manifold by *T*_min_.

All remaining triplet-derived states are collected into the interior triplet sector *T*_int_. If *T*_min_ exists, then *T*_int_ consists of the triplet-derived states with

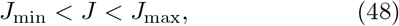

whereas if *T*_min_ does not exist it consists simply of the triplet-derived states with *J < J*_max_. The spin space is therefore partitioned naturally as

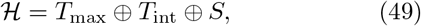

or, when the lower triplet-only manifold exists,

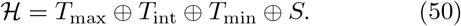

This is, however, a manifold-level classification. It identifies the *J* manifolds that are necessarily triplet-only by angular-momentum structure alone. It does not imply that non-extremal manifolds cannot contain additional pure-triplet zero-field eigenspaces. On the contrary, when further symmetries are present, especially those generated by equivalent nuclei, interior manifolds can separate into triplet-only and singlet-accessible zero-field eigenspaces. The detailed decomposition of *T*_int_ therefore depends on the specific hyperfine structure of the radical pair.

The correspondence with the simple one-nucleus model is immediate. For one spin 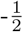 nucleus, 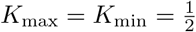 so 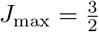 and no minimal triplet-only manifold exists. In that case

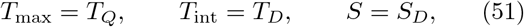

and the general pathway

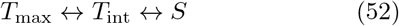

reduces exactly to

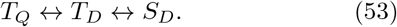

Table I illustrates the classification for a range of simple nuclear-spin sets.

**TABLE I.**
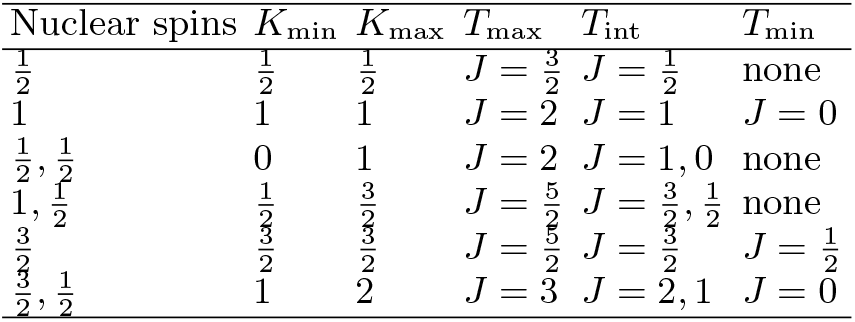
Examples of the maximal, interior, and minimal triplet manifolds for simple nuclear-spin sets. The nuclear spins listed are those present in the full radical pair.

In all of these cases, singlet-derived states occur only within the same non-extremal *J* manifolds listed under *T*_int_, whereas the maximal and, when present, minimal manifolds *T*_max_ and *T*_min_ are necessarily triplet-only at the manifold level.

For equal electron *g* values, the Zeeman interaction preserves total electron spin,

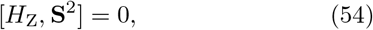

and therefore does not directly couple singlet- and triplet-derived states. Its direct action is purely within the triplet sector, where it mixes states belonging to different triplet manifolds. In particular, it can transfer amplitude from the maximal manifold *T*_max_ into the interior triplet sector *T*_int_, and, when present, also between *T*_int_ and the minimal manifold *T*_min_.

By contrast, the isotropic hyperfine interaction is responsible for communication between the interior triplet sector and the singlet sector within fixed *J, M* blocks. The essential LFE mechanism therefore remains the many-spin analogue of the one-nucleus pathway:

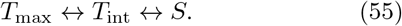

If a *T*_min_ manifold is present, it can participate in additional triplet-triplet redistribution, but it does not alter the basic sequential structure of the mechanism.

The singlet projector in the general problem is defined as,

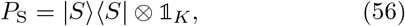

where 𝟙_*K*_ is the identity on the nuclear spin space. Thus *P*_S_ annihilates all states in *T*_max_ and, when present, all states in *T*_min_, while acting nontrivially only on singlet-containing components in the interior manifolds. This gives a simple physical summary: at zero field, part of the triplet manifold is isolated from the singlet-reactive sector because it is confined to triplet-only states; a weak magnetic field transfers amplitude from this triplet-only region into interior triplet states that share total-spin manifolds with singlet-derived states; and hyperfine coupling then transfers amplitude from those interior triplet states into the singlet sector.

The *T*_max_, *T*_int_, *T*_min_, and *S* classification developed here is exact for the isotropic spin Hamiltonian, where the zero-field problem retains full rotational symmetry and can therefore be organized by angular momentum. Additional interactions modify this picture to different degrees. Nuclear Zeeman terms are usually small in the weak-field regime and do not by themselves overturn the basic isotropic organisation, although they do lift residual degeneracies. By contrast, anisotropic hyperfine couplings and electron-electron dipolar interactions break rotational symmetry more fundamentally. In their presence, the total-spin labels used here are no longer exact quantum numbers, and the *T*_max_, *T*_int_, and *T*_min_ sectors cease to be exact symmetry-protected manifolds of the full Hamiltonian. The present framework should therefore be understood as an exact description of the isotropic problem, and as a useful reference point for analysing weaker anisotropic perturbations. Thus the *T*_max_, *T*_int_, *T*_min_ basis-state description leads to a simple intuitive picture of RP behaviour in zero and weak field. Figure 2 shows a diagrammatic representation of RP magnetic-field effects from zero field through weak field and into the regime where the Zeeman interaction dominates the hyperfine interaction. This picture encapsulates the RP behaviour without explicit reference to the details of the quantum states or coherences, providing a more intuitive representation of the RPM than existing formulations.

**FIG. 2.**
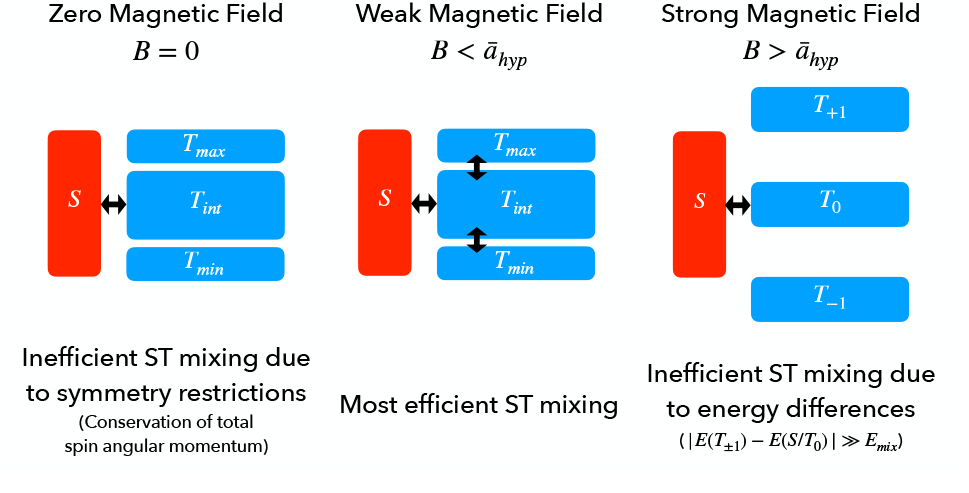
Intuitive picture of ST mixing from zero to high field.

## V. A PRACTICAL RECRUITMENT MEASURE FOR THE MAXIMUM LOW-FIELD EFFECT

The manifold classification above suggests a natural structural measure of the maximum LFE: the number of zero-field triplet-only states that are recruited into the singlet-accessible sector when a weak magnetic field is applied. This quantity is attractive because it measures, in a direct and physically transparent way, how much of the zero-field triplet-only state space is released into singlet-triplet communication when the field is turned on.

At the same time, a simple angular-momentum count is not sufficient. The decomposition into *T*_max_, *T*_int_, and *T*_min_ identifies which manifolds are triplet-only and which are singlet-accessible at the manifold level, but the actual number of states recruited at weak field also depends on the degeneracy structure of the zero-field Hamiltonian. Equivalent nuclei can create repeated degenerate blocks, and additional symmetry can produce symmetry-protected dark states. The recruitment measure must therefore be constructed from the exact zero-field eigenspaces rather than from manifold dimensions alone.

We therefore define

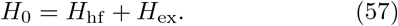

The eigenstates of *H*_0_ are then grouped into degenerate energy blocks. Let *P*_*E*_ denote the projector onto the zero-field eigenspace of energy *E*,

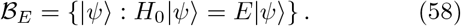

Restricting the singlet projector to this block gives

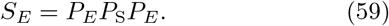

The kernel of *S*_*E*_ defines the triplet-only zero-field subspace within that block,

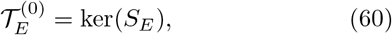

and the total dimension of the zero-field triplet-only sector is therefore

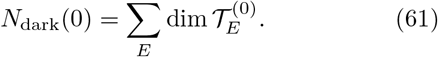

When a weak magnetic field is applied, the Hamiltonian becomes

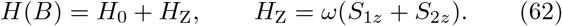

The same blockwise construction is then repeated for the eigenspaces of *H*(*B*), giving a field-dependent triplet-only dimension *N*_dark_(*B*). The number of triplet-only states recruited by the field is defined as

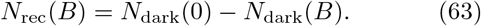

In practice, *N*_dark_(0) and *N*_dark_(*B*) are obtained by diagonalising the singlet projector within each energy-degenerate eigenspace and counting the number of zero eigenvalues to within a chosen numerical tolerance. This procedure is robust to arbitrary basis rotations within degenerate manifolds and naturally incorporates equivalent nuclei and other hyperfine symmetries.

Two associated normalised measures are then useful:

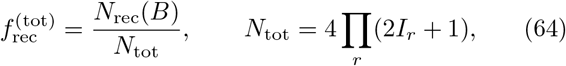

and

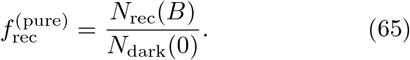

The first quantity measures the absolute size of the newly recruited state space relative to the whole Hilbert space. The second measures the fraction of the zero-field triplet-only sector that is released into the singlet-accessible sector. In what follows, however, the central quantity is the practical blockwise count *N*_rec_(*B*) itself.

These quantities should be interpreted as structural upper-bound indices for the LFE rather than as the LFE itself. The observed field effect will also depend on reaction rates, relaxation, exchange, and the amount of singlet character acquired by the recruited states.

The point of the recruitment measure is therefore not to replace a full dynamical calculation, but to isolate one of the key structural ingredients of the LFE. Specifically, *N*_rec_ quantifies how much of the zero-field triplet-dark state space becomes, in principle, connected to the singlet-reactive sector when a weak magnetic field is applied. In this sense it measures the size of the spin-space channel opened by the weak field. Radical pairs with larger recruitment counts possess a larger symmetry-allowed capacity for weak-field-induced singlet accessibility, whereas radical pairs with smaller recruitment counts possess less such capacity. The actually observed LFE can then be smaller, or strongly suppressed, depending on kinetics and other dynamical details. The recruitment measure should therefore be interpreted as an upper-bound indicator for the LFE rather than as a complete predictor of the final experimental signal.

The most obvious implication of the recruitment measure is that in general, the greater the number of hyperfine couplings and the larger the spin quantum numbers of the nuclei, the smaller the size of *T*_max_ (and where present *T*_min_) relative to *T*_*int*_, and thus the smaller the value of *f*_rec_. Therefore the one proton RP represents the case with the strongest possible LFE (and thus must be treated with caution when used for modelling real LFEs in general) and in general, greater numbers of spin-active nuclei serve to reduce the size of the LFE. Because at least one hyperfine coupling is needed to drive the LFE, the most effective RPs therefore have a dominant hyperfine coupling in one radical and as few as possible (and as weak as possible when present) couplings in the other radical.

The structural upper-bound indicator is important not only for identifying candidate RPs that may show large observable LFEs, but also for determining why, when no LFE is observed in an experiment, whether the reason is due to the structure of the RP itself or rather suppression by factors such as short RP lifetime or fast relaxation.

Simple Python code implementing the practical blockwise recruitment measure described above is provided in the supplementary material.

## VI. ILLUSTRATIVE TWO-PROTON EXAMPLES

The role of hyperfine symmetry is already evident in the four possible arrangements of two spin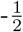 nuclei listed below. In all four cases the full Hilbert-space dimension is *N*_tot_ = 16, but the number of triplet-only zero-field eigenstates recruited by a weak field depends on how the hyperfine couplings are arranged:

1. two equivalent nuclei on radical 1,

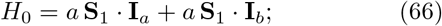
2. one nucleus on each radical, with equal couplings,

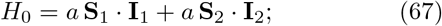
3. two nuclei on radical 1, with inequivalent couplings,

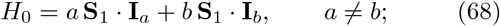
4. one nucleus on each radical, with inequivalent couplings,

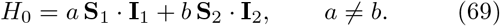

For the two inequivalent cases, explicit diagonalisation gives the generic result

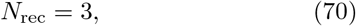

Physically, the weak field recruits the three non-stretched members (*M* = +1, 0, −1) of the maximal triplet manifold *T*_max_ into the singlet-accessible sector, while the stretched states remain triplet-only.

The two symmetric cases depart from this generic behaviour in different ways. For two equivalent nuclei on the same radical, the equivalence preserves an additional symmetry associated with the total nuclear spin of that radical. As a result, an extra triplet-only zero-field eigenstate appears within a non-extremal *J* manifold, giving

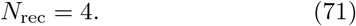

By contrast, when one equivalent nucleus is placed on each radical, the equality of the two couplings introduces a radical-exchange symmetry. In this case one of the nominally available weak-field channels becomes dark because the corresponding Zeeman matrix element vanishes by symmetry, leading to

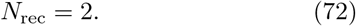

The results are summarised in Table II.

**TABLE II.** Weak-field recruitment counts *N*_rec_ for four arrangements of two spin 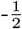 nuclei, obtained by blockwise dark-state counting.

| Case | Hyperfine arrangement | $N_{\text{rec}}$ |
| --- | --- | --- |
| 1 | two equivalent nuclei on radical 1 | 4 |
| 2 | one nucleus on each radical, equal couplings | 2 |
| 3 | two nuclei on radical 1, inequivalent couplings | 3 |
| 4 | one nucleus on each radical, inequivalent couplings | 3 |

These examples show that the recruitment count is not determined solely by the spin inventory of the radical pair. It also depends on the degeneracy structure and symmetry of the zero-field hyperfine Hamiltonian. Equivalent nuclei can therefore either increase or decrease the weak-field recruitment count relative to the generic inequivalent-coupling case.

In practice, the impact of the increase in *f*_rec_ in the case of equivalent nuclei in one radical is clearly observed as increase in the actual observed LFE in experiments^17^ and can now be simply rationalised in terms of the change in the recruitment measure.

## Supporting information

Python code and instructions

## Appendix A: Why the two symmetric two-proton cases behave differently

For completeness, we briefly show why the two symmetric two-proton arrangements give recruitment counts different from the generic value.

### 1. Two equivalent nuclei on radical 1

For case 1,

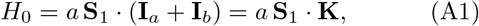

where

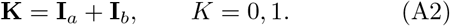

The hyperfine Hamiltonian is therefore diagonal in *K*. For *K* = 1, coupling 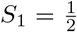 to *K* = 1 gives local resultant 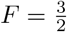 or 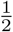, with energies

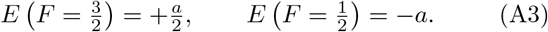

For *K* = 0, only 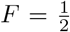 is possible, with energy *E* = 0. This sector is special because the nuclear pair is itself a singlet. The hyperfine interaction therefore vanishes there, and coupling to 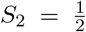 yields one pure-triplet *J* = 1 state and one pure-singlet *J* = 0 state.

The additional recruited state does not arise from that *K* = 0 sector. Instead, it appears within the *K* = 1, 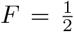 manifold at energy −*a*, where there occur both a *J* = 1 singlet-accessible block and a *J* = 0 state that is necessarily triplet-only. Because this *J* = 0 state can arise only from electron-triplet coupling when *K* = 1, it is a symmetry-protected triplet-only zero-field eigenstate. It is degenerate with the corresponding *J* = 1 block and is connected to its *M* = 0 component by the Zeeman perturbation. Thus the three non-stretched members of the *J* = 2 manifold are recruited, together with one additional symmetry-protected triplet-only state, giving a total recruitment count of four.

### 2. One nucleus on each radical with equal couplings

For case 2,

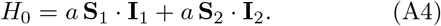

Each radical may be in a local singlet (*F*_*i*_ = 0) or triplet (*F*_*i*_ = 1) state. The two *J* = 1 sectors arising from (*F*_1_, *F*_2_) = (1, 0) and (0, 1) are exactly degenerate when the couplings are equal and can therefore be reorganised into symmetry-adapted combinations under interchange of the two radicals. One such combination is triplet-only, while another is singlet-accessible.

The Zeeman operator

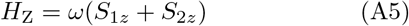

is symmetric under interchange of the two radicals. As a consequence, the *M* = 0 matrix element between the triplet-only and singlet-accessible combinations vanishes identically, whereas the corresponding *M* = ±1 matrix elements are nonzero. The *M* = 0 channel is therefore dark. Only the two *M* = ± 1 directions are recruited, giving a total count of two.

Taken together, these two examples show the two main ways in which symmetry modifies the generic recruitment count. In one case, symmetry creates an additional triplet-only zero-field eigenspace that can couple into the singlet-accessible sector. In the other, symmetry creates a dark combination for which the Zeeman coupling vanishes.

## SUPPLEMENTARY MATERIAL

Supplementary material provides simple Python code implementing the practical blockwise recruitment measure introduced in Sec. V.

## ACKNOWLEDGMENTS

The author thanks Noboru Ikeya for helpful discussions. This work was supported by Japan Society for the Promotion of Science (JSPS) Grants-in-Aid for Scientific Research (Grant No. 23K26612).

