## Supplementary material for "The low-field effect in radical pairs: a zero-field singlet-triplet basis picture": Python code and instructions: rp_recruitment.py

```
"""Supplementary-information code for the recruitment measure.

This script implements the practical weak-field recruitment count used in the
manuscript for isotropic radical-pair spin Hamiltonians. For a radical pair
with arbitrary isotropic hyperfine couplings and optional isotropic exchange,
it computes

    N_rec(B) = N_dark(0) - N_dark(B),

where N_dark is the dimension of the triplet-only (dark) subspace obtained by
diagonalizing the singlet projector within each energy-degenerate eigenspace
of the Hamiltonian.

The implementation is written for clarity and transparency rather than maximal
performance. It is intended for supporting-information calculations on
moderate Hilbert-space dimensions and reproduces the benchmark two-proton
examples discussed in the manuscript.
"""

from __future__ import annotations

from dataclasses import dataclass
from typing import Any, List, Sequence

import numpy as np


Array = Any


@dataclass(frozen=True)
class Nucleus:
    spin: float
    coupling: float


@dataclass(frozen=True)
class RadicalPairModel:
    radical1: Sequence[Nucleus]
    radical2: Sequence[Nucleus]
    exchange: float = 0.0  # H_ex = 2 J_ex S1.S2


@dataclass(frozen=True)
class RecruitmentMeasures:
    n_total: int
    n_dark_zero: int
    n_dark_weak: int
    n_rec: int
    f_rec_total: float
    f_rec_pure: float


def spin_matrices(spin: float) -> tuple[Array, Array, Array, Array]:
    two_s = round(2 * spin)
    if abs(two_s - 2 * spin) > 1e-9 or spin < 0:
        raise ValueError(f"Invalid spin quantum number: {spin}")

    dim = two_s + 1
    m_vals = np.arange(spin, -spin - 1, -1, dtype=float)

    s_plus = np.zeros((dim, dim), dtype=complex)
    s_minus = np.zeros((dim, dim), dtype=complex)
    for i, m in enumerate(m_vals[:-1]):
        coeff = np.sqrt(spin * (spin + 1) - m * (m - 1))
        s_minus[i + 1, i] = coeff
        s_plus[i, i + 1] = coeff

    sx = 0.5 * (s_plus + s_minus)
    sy = -0.5j * (s_plus - s_minus)
    sz = np.diag(m_vals)
    ident = np.eye(dim, dtype=complex)
    return sx, sy, sz, ident


def kron_all(operators: Sequence[Array]) -> Array:
    result = operators[0]
    for op in operators[1:]:
        result = np.kron(result, op)
    return result


def embed_single_site(
    site_ops: Sequence[tuple[Array, Array, Array, Array]], index: int, component: int
) -> Array:
    ops: List[Array] = []
    for i, (sx, sy, sz, ident) in enumerate(site_ops):
        if i == index:
            ops.append((sx, sy, sz)[component])
        else:
            ops.append(ident)
    return kron_all(ops)


def group_indices(values: Array, tol: float) -> List[List[int]]:
    groups: List[List[int]] = []
    current: List[int] = [0]
    for i in range(1, len(values)):
        if abs(values[i] - values[current[0]]) <= tol:
            current.append(i)
        else:
            groups.append(current)
            current = [i]
    groups.append(current)
    return groups


def dot_operator_sum(left_ops: Sequence[Array], right_ops: Sequence[Array]) -> Array:
    """Return sum_k A_k B_k without relying on Python's scalar-start sum."""

    result = np.zeros_like(left_ops[0])
    for left, right in zip(left_ops, right_ops):
        # `einsum` avoids occasional backend-specific matmul warnings seen in
        # some GUI Python environments while producing the same dense product.
        result += np.einsum("ij,jk->ik", left, right, optimize=True)
    return result


class RadicalPairRecruitment:
    def __init__(self, model: RadicalPairModel):
        self.model = model
        self._build_operators()

    def _build_operators(self) -> None:
        nuclear_sites = list(self.model.radical1) + list(self.model.radical2)
        self._rad1_nuclear_indices = list(range(2, 2 + len(self.model.radical1)))
        self._rad2_nuclear_indices = list(
            range(2 + len(self.model.radical1), 2 + len(nuclear_sites))
        )

        site_ops: List[tuple[Array, Array, Array, Array]] = [
            spin_matrices(0.5),
            spin_matrices(0.5),
        ]
        for nucleus in nuclear_sites:
            site_ops.append(spin_matrices(nucleus.spin))

        self.site_ops = site_ops
        self.dimension = int(np.prod([ops[3].shape[0] for ops in site_ops]))

        self.s1 = tuple(embed_single_site(site_ops, 0, comp) for comp in range(3))
        self.s2 = tuple(embed_single_site(site_ops, 1, comp) for comp in range(3))
        self.nuclear_ops = [
            tuple(embed_single_site(site_ops, i, comp) for comp in range(3))
            for i in range(2, len(site_ops))
        ]

        self.p_s = self._build_singlet_projector()
        self.h0 = self._build_zero_field_hamiltonian()

    def _build_singlet_projector(self) -> Array:
        singlet = np.array(
            [0.0, 1.0 / np.sqrt(2.0), -1.0 / np.sqrt(2.0), 0.0], dtype=complex
        )
        p_s_e = np.outer(singlet, singlet.conj())
        nuclear_dim = self.dimension // 4
        return np.kron(p_s_e, np.eye(nuclear_dim, dtype=complex))

    def _build_zero_field_hamiltonian(self) -> Array:
        h = np.zeros((self.dimension, self.dimension), dtype=complex)

        for idx, nucleus in enumerate(self.model.radical1):
            i_ops = self.nuclear_ops[idx]
            h = h + nucleus.coupling * dot_operator_sum(self.s1, i_ops)

        offset = len(self.model.radical1)
        for idx, nucleus in enumerate(self.model.radical2):
            i_ops = self.nuclear_ops[offset + idx]
            h = h + nucleus.coupling * dot_operator_sum(self.s2, i_ops)

        if abs(self.model.exchange) > 0.0:
            h = h + 2.0 * self.model.exchange * dot_operator_sum(self.s1, self.s2)

        return h

    def zeeman_hamiltonian(self, omega: float) -> Array:
        return omega * (self.s1[2] + self.s2[2])

    def dark_state_count_at_field(
        self,
        omega: float,
        energy_tol: float = 1e-9,
        singlet_tol: float = 1e-9,
    ) -> int:
        h = self.h0 + self.zeeman_hamiltonian(omega)
        return self._dark_state_dimension_of_hamiltonian(
            h, energy_tol=energy_tol, singlet_tol=singlet_tol
        )

    def _dark_state_dimension_of_hamiltonian(
        self,
        hamiltonian: Array,
        energy_tol: float = 1e-9,
        singlet_tol: float = 1e-9,
    ) -> int:
        evals, vecs = np.linalg.eigh(hamiltonian)
        energy_groups = group_indices(evals, energy_tol)
        dark_dim_total = 0

        for e_group in energy_groups:
            v_e = vecs[:, e_group]
            s_e = v_e.conj().T @ self.p_s @ v_e
            s_vals = np.linalg.eigvalsh(s_e)
            dark_dim_total += int(np.sum(s_vals <= singlet_tol))

        return dark_dim_total

    def recruitment_measures(
        self,
        omega_weak: float,
        energy_tol: float = 1e-9,
        singlet_tol: float = 1e-9,
    ) -> RecruitmentMeasures:
        n_dark_zero = self.dark_state_count_at_field(
            omega=0.0, energy_tol=energy_tol, singlet_tol=singlet_tol
        )
        n_dark_weak = self.dark_state_count_at_field(
            omega_weak, energy_tol=energy_tol, singlet_tol=singlet_tol
        )
        n_rec = n_dark_zero - n_dark_weak

        n_total = self.dimension
        f_rec_total = n_rec / n_total
        f_rec_pure = n_rec / n_dark_zero if n_dark_zero else 0.0

        return RecruitmentMeasures(
            n_total=n_total,
            n_dark_zero=n_dark_zero,
            n_dark_weak=n_dark_weak,
            n_rec=n_rec,
            f_rec_total=f_rec_total,
            f_rec_pure=f_rec_pure,
        )


def example_models(a: float = 1.0, b: float = 0.7) -> dict[str, RadicalPairModel]:
    return {
        "case_1_equiv_same_radical": RadicalPairModel(
            radical1=[Nucleus(0.5, a), Nucleus(0.5, a)],
            radical2=[],
        ),
        "case_2_equiv_split_radicals": RadicalPairModel(
            radical1=[Nucleus(0.5, a)],
            radical2=[Nucleus(0.5, a)],
        ),
        "case_3_inequiv_same_radical": RadicalPairModel(
            radical1=[Nucleus(0.5, a), Nucleus(0.5, b)],
            radical2=[],
        ),
        "case_4_inequiv_split_radicals": RadicalPairModel(
            radical1=[Nucleus(0.5, a)],
            radical2=[Nucleus(0.5, b)],
        ),
    }


def print_summary(name: str, measures: RecruitmentMeasures) -> None:
    print(name)
    print(f"  N_total        = {measures.n_total}")
    print(f"  N_dark(0)      = {measures.n_dark_zero}")
    print(f"  N_dark(weak)   = {measures.n_dark_weak}")
    print(f"  N_rec          = {measures.n_rec}")
    print(f"  f_rec^(tot)    = {measures.f_rec_total:.6f}")
    print(f"  f_rec^(pure)   = {measures.f_rec_pure:.6f}")
    print()


if __name__ == "__main__":
    omega_weak = 1.0e-4
    for name, model in example_models().items():
        calculator = RadicalPairRecruitment(model)
        result = calculator.recruitment_measures(omega_weak=omega_weak)
        print_summary(name, result)
```
