## Supplementary material for "The low-field effect in radical pairs: a zero-field singlet-triplet basis picture": Python code and instructions: rp_recruitment_README.txt

Supplementary-information code for the recruitment measure

File included

`rp_recruitment.py` provides a Python implementation of the practical weak-field recruitment measure for isotropic radical-pair spin Hamiltonians.

Purpose

The script evaluates the quantity

`N_rec(B) = N_dark(0) - N_dark(B)`,

where `N_dark` is the dimension of the triplet-only (dark) subspace obtained by diagonalizing the singlet projector within each energy-degenerate eigenspace of the Hamiltonian. It also reports the normalized quantities `f_rec^(tot)` and `f_rec^(pure)` used in the manuscript.

Method

For a chosen magnetic field, the script:

1. diagonalizes the Hamiltonian,
2. groups the eigenvectors into energy-degenerate blocks,
3. diagonalizes the singlet projector `P_S` within each block,
4. counts the zero eigenvalues of the restricted projector to obtain the number of dark states in that block,
5. sums over all blocks to obtain `N_dark`.

This blockwise procedure is essential because individual eigenvectors within a degenerate eigenspace are not unique. Counting dark states blockwise makes the result basis-independent and correctly handles equivalent nuclei and other hyperfine symmetries.

Model assumptions

- Two electron spins, each with `S = 1/2`
- Arbitrary isotropic hyperfine couplings to nuclei on radical 1 and radical 2
- Optional isotropic exchange with convention `H_ex = 2 J_ex S1.S2`
- Equal electron `g` values for the Zeeman term
- No nuclear Zeeman, dipolar, anisotropic hyperfine, or `Delta g` terms

Model specification

Each nucleus is defined as `Nucleus(spin, coupling)`. A radical pair is defined as `RadicalPairModel(radical1=[...], radical2=[...], exchange=...)`.

Illustrative use

```python
from rp_recruitment import Nucleus, RadicalPairModel, RadicalPairRecruitment

model = RadicalPairModel(
    radical1=[Nucleus(0.5, 1.0), Nucleus(0.5, 0.7)],
    radical2=[Nucleus(0.5, 0.4)],
    exchange=0.0,
)

calculator = RadicalPairRecruitment(model)
measures = calculator.recruitment_measures(omega_weak=1.0e-4)

print(measures.n_dark_zero)
print(measures.n_dark_weak)
print(measures.n_rec)
```

Benchmark examples included in the script

- two equivalent spin-1/2 nuclei on one radical
- one spin-1/2 nucleus on each radical with equal couplings
- two spin-1/2 nuclei on one radical with unequal couplings
- one spin-1/2 nucleus on each radical with unequal couplings

Dependency

The script requires `numpy`.

General note

The implementation is written for clarity and transparency rather than large-scale optimization. For larger spin systems, sparse or symmetry-adapted implementations may be preferable.
